# The expression of class II major histocompatibility molecules on breast tumors delays T cell exhaustion, expands the T cell repertoire and slows tumor growth

**DOI:** 10.1101/294124

**Authors:** Tyler R. McCaw, Mei Li, Dmytro Starenki, Sara J. Cooper, Selene Meza-Perez, Rebecca C. Arend, Albert F. LoBuglio, Donald J. Buchsbaum, Andres Forero, Troy D. Randall

## Abstract

The expression of major histocompatibility complex II (MHCII) on tumor cells correlates with survival and responsiveness to immunotherapy. However, the mechanisms underlying these observations are poorly defined. Using a murine breast tumor line, we tested how MHCII expression affected anti-tumor immunity. We found that MHCII-expressing tumors grew more slowly than controls and recruited more functional CD4+ and CD8+ T cells. Additionally, MHCII-expressing tumors contained more TCR clonotypes expanded to a larger degree than control tumors. Functional CD8+ T cells in tumors depended on CD4+ T cells. However, both CD4+ and CD8+ T cells eventually became exhausted, even in MHCII-expressing tumors. PD1 blockade had no impact on tumor growth, potentially because tumor cells poorly expressed PD-L1. These results suggest tumor cell expression of MHCII facilitates the local activation of CD4+ T cells and indirectly helps the activation and expansion of CD8+ T cells, but by itself, cannot prevent T cell exhaustion.

**Précis:** The expression of MHCII on tumor cells augments CD4 and CD8 T cell responses, expands the TCR repertoire and delays exhaustion. Hence, strategies to induce MHCII expression may be a powerful adjuvant to immunotherapeutic regimens of solid tumors.

## INTRODUCTION

The capacity of the adaptive immune system to spontaneously recognize and respond to tumors is now generally accepted (1, 2), although these responses are often counteracted by a variety of local immune-inhibitory mechanisms, including signaling through T cell inhibitory receptors (3), such as PD-1. In fact, the therapeutic inhibition of these receptors with antibodies, known as checkpoint inhibitors, can elicit remarkable clinical responses, especially in melanoma and non-small cell lung cancer (4, 5). However, for many tumor types, only a minority of patients experiences a therapeutic benefit, a phenomenon ascribed to variations in the tumor microenvironment as well as the magnitude of the local immune response prior to treatment (6, 7). Thus, it is important to understand how variations in gene expression by tumor cells and the tumor microenvironment contribute to clinical outcomes.

The expression of major histocompatibility class II (MHCII) molecules by tumor cells is one variable that strongly predicts clinical outcomes. Although the expression of MHCII is normally restricted to professional antigen-presenting cells, such as dendritic cells, macrophages and B cells, our group and others have shown that aberrant expression of MHCII pathway components on melanoma (8), colorectal carcinoma (9, 10), Hodgkin’s lymphoma (11), diffuse large B cell lymphoma (12), ovarian cancer (13), and breast cancer (14, 15), is consistently associated with both improved patient outcomes and more vigorous immune responses. In fact, we demonstrated that coordinated expression of 13 MHCII pathway genes in samples of triple negative breast cancer (TNBC) is positively associated with prolonged progression free survival and the accumulation of tumor-infiltrating lymphocytes (TILs) (14). Even more strikingly, the expression of the class II transcriptional activator (CIITA), the master regulator of MHCII expression (16), is independently associated with progression free survival (14). These data suggest that the ability of tumor cells to present antigen to CD4+ T cells is a critical component of functional anti-tumor immune responses.

Unlike CD8+ T cells, CD4+ T cells are typically not cytolytic, but instead make cytokines that promote inflammation and cellular activation. Moreover, CD4+ T cells provide “help” to CD8+ T cells by promoting dendritic cell activation and cross-presentation (17, 18). Thus, the expression of MHCII on tumor cells likely promotes the local activation of CD4+ T cells, which indirectly help CD8+ T cells kill tumor cells. Consistent with this idea, the ectopic expression of MHCII on murine tumor cells promotes the local expression of IFNγ, adaptive immunity and tumor rejection in the context of mammary adenocarcinoma (19, 20). Rejection of MHCII-expressing tumors occurs independently of B cells, dendritic cells, macrophages, and neutrophils, but instead relies on CD4+ and CD8+ T cells as well as the production of IFNγ (20). However, none of these studies examined how MHCII expression by tumor cells altered the functional abilities of either CD4+ or CD8+ T cells.

T cells responding to persistent antigens, such as those expressed during chronic infections or by tumor cells, often become exhausted (21). Exhausted T cells are unable to express cytokines or kill target cells (22), a phenotype that allows tumor outgrowth despite initial immune activation. T cell exhaustion is caused by chronic TCR stimulation, often in the absence of costimulation or cytokines (21). For example, CD8+ T cells that fail to receive help from CD4+ T cells acquire a “helpless” or exhausted phenotype (23, 24). Inhibitory receptors on T cells, such as PD-1, also contribute to T cell exhaustion (25), particularly when their ligands are expressed on target cells, such as tumor cells. Interestingly, PD-1 blockade is more successful in melanoma patients whose tumors also express MHCII (8). Thus, MHCII expression on tumor cells may promote successful anti-tumor immunity by enhancing CD4+ helper functions and preventing the exhaustion of CD8+ T cells or allowing exhausted T cells to recover when inhibitory molecules are blocked.

Here we tested how ectopic expression of MHCII on murine TS/A breast tumor cells altered tumor-specific CD4+ and CD8+ T cell responses. We found that local MHCII expression increased the magnitude and duration of both CD4+ and CD8+ T cell responses in tumors. We also observed alterations in the T cell repertoire, with more clonotypes expanded to a larger extent in MHCII-expressing tumors. Interestingly, although effector functions of T cells were significantly augmented and prolonged in MHCII-expressing tumors, both CD4+ and CD8+ T cells eventually became exhausted. Moreover, while more CD8+ T cells in MHCII-expressing tumors expressed PD-1 and higher amounts thereof, the blockade of PD-1 did not reverse the exhausted phenotype of either CD4+ or CD8+ T cells or impact tumor growth. These data suggest that MHCII expression on tumor cells expands the T cell repertoire and delays, but does not prevent, T cell exhaustion, which temporarily abates tumor growth. However, additional suppressive mechanisms must be overcome to achieve tumor eradication.

## MATERIALS and METHODS

### Cell culture, transfection, and selection

The TS/A murine mammary adenocarcinoma cell line was generously provided by Dr. Roberto S. Accolla, Department of Clinical and Biological Sciences, University of Insubria, Italy. TS/A cells were obtained at passage 22 and passaged 2 times prior to freezing. Cells were authenticated by assessing MHC haplotype via flow cytometry and by detection of antigens from murine leukemia virus. Transfected and control TS/A cells were also confirmed by gene expression of MHCII pathway gene products using nanostring assay. Frozen aliquots from passage 24 were thawed and expanded prior to injection. Cells were cultured in DMEM (Corning) supplemented with 10% fetal bovine serum (FBS; HyClone Laboratories, Inc.), and passaged by dissociating confluent monolayers with 0.05% trypsin, 0.53 mM EDTA without sodium bicarbonate (Corning), washing and reculturing. Cells were transfected with pcDNA3.1 with or without the hCIITA cDNA using the Xfect Transfection Reagent (Clontech) and selected in 500 µg/mL geneticin (Life Technologies). Clones that stably expressed MHCII were further selected by cell sorting. The expression of hCIITA, MHCII and CD74 in transfected cells was confirmed by Western blot using antibodies against hCIITA (E-12, Santa Cruz Biotechnology), MHCII (M5/114, Millipore), and CD74 (R&D Systems). Cells tested negative for mycoplasma (and 13 other mouse pathogens) via PCR performed by Charles River Research Animal Diagnostic Services on June 2015.

### RNA isolation and nanostring analysis

Total RNA was purified from excised tumors using the RNeasy Mini kit (Qiagen) and quantified by optical density at 260 nm using a DeNovix DS-11 spectrophotometer (DeNovix Inc.). Total RNA from blood was obtained from ∼1 mL (from mouse cardiac puncture) of whole blood collected into Paxgene RNA tubes using the Paxgene blood RNA kit (Qiagen) according to manufacturer instructions. The quality of RNA was assessed using Bioanalyzer 2100 (Agilent) and final concentration determined by Qubit (Life Technologies).

For nanostring analysis, 100 ng of purified RNA was added to 3 μL of Reporter CodeSet and 2 μL Capture ProbeSet in accordance with manufacturer’s recommendations (NanoString Technologies), and hybridized at 65°C overnight. Samples were processed the following day using the nCounter DxPrep Station and nCounter Dx Digital Analyzer (NanoString Technologies). Consumables were provided as an nCounter master kit by the manufacturer (NanoString Technologies). Data was normalized to housekeeping genes and analyzed using the nSolver 2.6 software. Heat maps were created using Cluster 3.0 and Java TreeView-1.1.6r4.

### Mice, tumor administration and antibody treatments

BALB/c mice of at least 6 weeks of age were purchased from Charles River Laboratories International, Inc. BALB/c.scid mice (CBySmn.CB17-*Prkdc*^*scid*^/J) were purchased from The Jackson Laboratory. Each experiment or time point represents cohorts of n=5 done at least in duplicate. Mice were injected with 1 × 10^5^ TS/A or TS/A-hCIITA tumor cells into the mammary fat pad on day 0. The longest (length, L) and shortest (width, W) tumor dimensions were measured by caliper and tumor volume was calculated using the formula, 0.4 x L x W^2^. For depletion experiments, 200 μg of CD4-depleting antibody (clone GK1.5, BioXCell) or CD25-depleting antibody (clone PC-61.5.3, BioXCell) or isotype-matched control antibody (clone LTF-2 or HRPN, respectively; BioXCell) were administered prior to tumor cell injection (day −1) and again 3 days following injection. For PD-1 blockade, 200 µg of anti-PD-1 antibody (clone J43, BioXCell) or isotype-matched control antibody (Armenian Hamster IgG, BioXCell) was administered on days 8, 11, and 14 post tumor cell injection. All procedures involving animals were approved by the University of Alabama at Birmingham Institutional Animal Care and Use Committee (IACUC) in protocol 09854.

### Tumor disassociation and T cell restimulation

Tumors were excised, weighed using an AL54 analytical balance (Mettler Toledo), and diced using a scalpel. Tumor fragments were then incubated in 2 mL RPMI1640 media (Lonza) supplemented with 5% FBS, 1.25 mg collagenase (c7657, Sigma) and 150 U DNase (d5025, Sigma), followed by gentle shaking at 200 rpm for 35 minutes at 37°C. Cell suspensions were filtered through 70 µm nylon cell strainers (Corning) and used for flow cytometry or T cell restimulation. For T cell stimulation, cells were washed and resuspended in RPMI1640, 5% FBS, 5 ng/mL phorbol 12-myristate 13-acetate (Sigma), 65 ng/mL ionomycin (ThermoFisher Scientific), and 10 µg/mL brefeldin A (Sigma) and cultured at 37°C for 5 hours.

### Flow cytometry and antibodies

Cell cycle analysis was performed by releasing non-confluent TS/A and TS/A-hCIITA cells from culture dishes using Accutase with 0.5 mM EDTA (Innovative Cell Technologies), fixing with 70% ethanol at 4°C overnight, and incubating them with 50 ug/ml propidium iodide (ThermoFisher Scientific) in PBS, 0.1% Triton-X100 (Sigma), and 50 ug/ml RNase at 37°C for 20 minutes in the dark. DNA content was determined using a BD FACSCalibur (BD Biosystems) flow cytometer and analyzed by ModFit LT version 3.3 (Verity Software House).

For immunophenotyping, cell suspensions from disassociated tumors were washed and resuspended in staining solution-PBS with 2% donor calf serum and 10 μg/ml FcBlock (2.4G2 -BioXCell)-for 10 min on ice before staining with fluorochrome-conjugated antibodies. Extracellular staining was done for 30 minutes, following by washing and fixation with 10% neutral buffered formalin solution (Sigma). Intracellular staining was performed on fixed cells permeablized with 0.1% IGEPAL CO-630 (Sigma) made in staining solution for 45 minutes. All staining was performed in the dark at 4°C. Antibodies against I-A/I-E (M5/114.15.2) and CD4 (GK1.5) were obtained from BioLegend). Antibodies against IFNγ (XMG1.2) and CD8 (53-6.7) were obtained from BD Biosciences. Antibodies against CD3 (17A2), KLRG1 (2F1), Granzyme B (NGZB), CD45.2 (104) and PD-1 (J43), were obtained from eBioscience. The MuLV class I tetramer-H2L^d^ (SPSYVYHQF) was obtained from the NIH Tetramer Core Facility at Emory University. The LIVE/DEAD red fixable dye was obtained from Life Technologies. Samples were run on a BDFACS Canto II system (BD Biosciences) data was analyzed using FlowJo version 9.9.

### T cell repertoire sequencing and analysis

For TCR library preparation, we used the commercially available iRepertoire platform (iRepertoire, Huntsville, AL) for nested amplicon arm-PCR of the CDR3 of the mouse TCR β-chains and addition of adaptors for Illumina platform sequencing. Reverse transcription of 300 to 500 ng of RNA was conducted with a One-Step reverse transcription and amplification kit (Qiagen) according to the manufacturer’s protocol. The PCR product was purified using Ampure XP magnetic beads (Agencourt), and secondary amplification of the resulting product was performed (TopTaq PCR Kit, Qiagen), allowing addition of Illumina adapter sequences (manufacturer’s protocol). Libraries were purified with Ampure XP magnetic beads and sequenced using Illumina MiSeq 150 nt paired-end read-length. The TCR CDR3 sequences were extracted from the raw sequencing data by iRepertoire. Briefly, raw paired-end fastq files were first demultiplexed based on the internal 6-nt barcode sequences added during library construction. The merged reads were mapped using a Smith-Waterman algorithm to germline V, D, J and C reference sequences downloaded from the IMGT web site (http://www.imgt.org). To define the CDR3 region, the position of CDR3 boundaries of reference sequences from the IMGT database were migrated onto reads through mapping results and the resulting CDR3 regions were extracted and translated into amino acids. Reading frames amino acid motif that was uninterrupted by a stop codon were identified as productive CDR3 amino acid sequences. Reads were filtered and error-corrected using iRepertoire proprietary SMART algorithm.

### Statistical Analysis

Hypothesis of difference testing between pooled replicate group means at each time point was done using multiple, unpaired independent t-tests with the Holm-Sidak method and assuming α=0.05. Statistical analyses were calculated using GraphPad Prism version 7.0a and all reported p-values are two-tailed. The TCR high-throughput sequencing data were analyzed in R environment using tcR package and common R routines. Diversity was measured using D50 immune repertoire diversity index. CDR3 sequence similarity between repertoires was assessed using overlap index and weighted Horn’s index available in the tcR package.

## RESULTS

### Transfection with hCIITA promotes MHCII expression on TS/A breast cancer cells

Our previous studies show that patients whose triple negative breast tumors expressed MHCII have greater numbers of tumor infiltrating lymphocytes (TILs) and experience prolonged progression free survival relative to those lacking MHCII expression (14). Given the central role of MHCII in stimulating CD4+ T cell responses to antigen, we sought to define the contribution of tumor-specific MHCII expression in the recruitment or local activation of TILs. To achieve this goal, we transfected TS/A cells, a murine mammary adenocarcinoma cell line, with the human class II transcriptional activator (hCIITA) or with empty vector. By Western blot analysis, we found that TS/A cells transfected with hCIITA expressed hCIITA as well as murine CD74 (invariant chain) and MHCII (Figure 1A), whereas cells transfected with empty vector did not. Critically, we found that hCIITA-transfected cells also expressed high levels of murine MHCII on the cell surface (Figure 1B). To assess whether hCIITA expression altered cellular proliferation, we used propidium iodide to measure DNA content and found that the cell cycling times of hCIITA-expressing and control TS/A cells were nearly equivalent (Figure 1C). These data demonstrate that hCIITA promotes the expression of genes in the MHCII pathway and drives murine MHCII expression on the cell surface of TS/A breast cancer cells.

**Figure 1.**
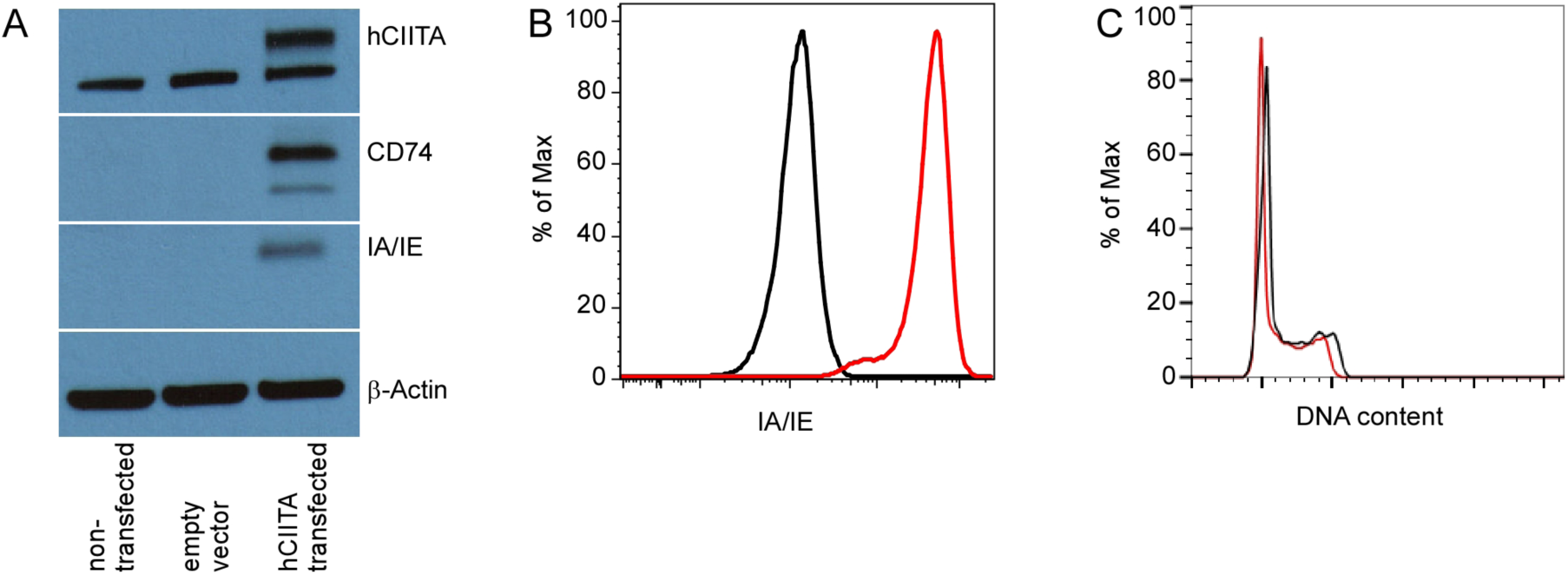
Transfection with hCIITA drives surface expression of murine MHCII. A. Protein expression of hCIITA, CD74, and IA/IE was detected by Western Blot on TS/A cells or TS/A cells transfected with hCIITA or empty vector. B. MHCII surface expression on TS/A control (black histogram) or TS/A-hCIITA (red histogram) cells was demonstrated by flow cytometry. C. DNA content of TS/A control (black histogram) or TS/A-hCIITA (red histogram) cells was determined by propidium iodide staining. These procedures were performed twice with similar results.

### Surface expression of MHCII impairs tumor growth *in vivo*

To test whether MHCII expression was maintained on CIITA-transfected TS/A cells *in vivo*, we injected TS/A and TS/A-hCIITA cells into the mammary fat pads of BALB/c mice and used flow cytometry to assess the expression of MHCII on CD45^neg^ cells from disassociated tumor masses 14 days later. Similar to our findings *in vitro*, we observed that the majority of CD45^neg^ cells from TS/A-hCIITA tumors expressed high levels of MHCII (Figure 2A), whereas CD45^neg^ cells from control tumors did not. However, in contrast to their similar growth kinetics *in vitro*, we found that TS/A-hCIITA tumors grew more slowly than control TS/A tumors, as measured by tumor volume (Figure 2B) and excised tumor mass (Figure 2C). To test whether this difference in tumor growth was dependent on adaptive immunity, we injected tumor cells into BALB/c.scid mice and found that the growth of TS/A-CIITA tumors was indistinguishable from the growth of control TS/A tumors over the same period (Figure 2D). These results suggested that the observed difference in growth of MHCII-positive and MHCII-negative tumors in wild type mice is mediated by an adaptive immune response.

**Figure 2.**
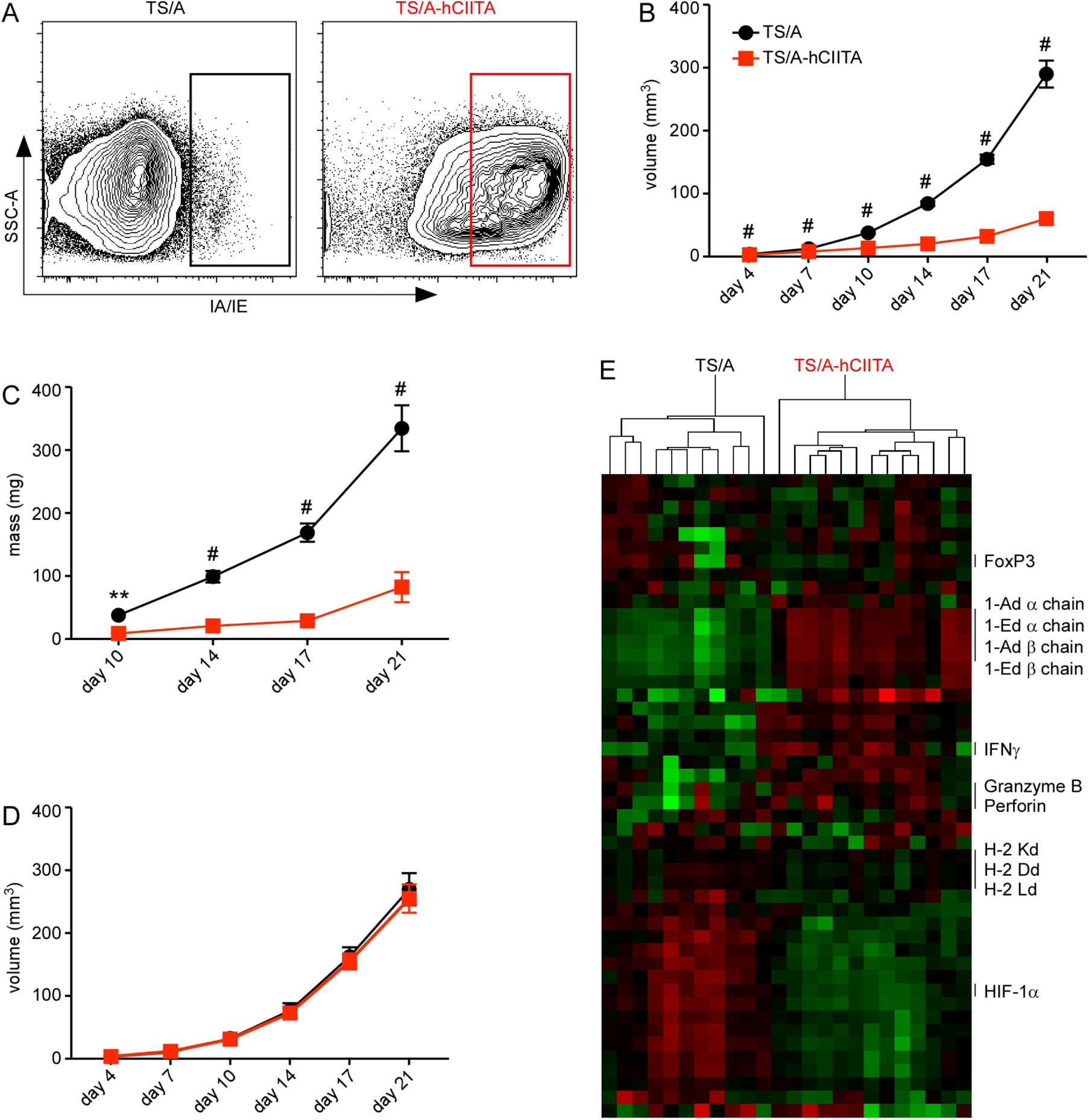
MHCII expression on tumor cells impairs tumor growth via adaptive immunity. TS/A and TS/A-hCIITA cells were injected into the mammary fat pad of BALB/c mice. A. MHCII surface expression on live, CD45^neg^ cells from disassociated tumors was measured by flow cytometry on day 14. B. Tumor volume was monitored over time in at least 20 mice per group per timepoint. C. Tumor mass was determined in 10 samples per group per timepoint. D. TS/A and TS/A-hCIITA cells were injected into the mammary fat pad of BALB/c-scid mice and tumor volume was monitored over time in 10 mice per group. E. TS/A (black) and TS/A-hCIITA (red) tumors were excised on days 10, 14, 17, and 21 after implantation, RNA was extracted and analyzed by Nanostring assay. Heat map shows fold increases in red and fold decreases in green. Error bars represent standard error of the mean. Statistical difference is expressed as *p<0.05, **p<0.005, ^#^p<0.0005.

To better understand how expression of the MHCII pathway in tumor cells impacted the local microenvironment, we excised whole tumors from recipient BALB/c mice on days 10, 14, 17, and 21, prepared mRNA and analyzed the expression of a panel of 48 genes using a NanoString assay. As expected, we found that reads of mRNAs encoding MHCII α and β chains were more numerous in TS/A-hCIITA tumors relative to control tumors across time points (Figure 2E), whereas reads of mRNAs encoding MHC class I molecules were largely similar. We also observed that several mediators of immunity and cytotoxicity - IFNγ, granzyme B and perforin - were upregulated in TS/A-hCIITA tumors relative to control tumors (Figure 2E). Interestingly, we found that the master transcription factor for regulatory T cells, FoxP3, was expressed similarly in TS/A-hCIITA and control tumors. In contrast, we found that the expression of other genes, such as HIF-1a, a transcription factor that facilitates tumor cell survival in hypoxic conditions (26) and promotes the ability of tumor-associated macrophages to suppress T cells (27), was consistently reduced in TS/A-hCIITA tumors. Together, these data demonstrated that MHCII expression on TS/A tumors promoted adaptive immune responses, altered the tumor microenvironment, and impaired tumor growth.

### MHCII-expressing tumors have increased CD4+ T cell infiltration and activation

Given that MHCII molecules stimulate CD4+ T cells, we hypothesized that MHCII-expressing tumor cells would bolster the local CD4+ T cell response. To test this possibility, we harvested TS/A-hCIITA and control tumors on days 10, 14, 17, and 21 after injection, restimulated the infiltrating cells *ex vivo* for 5 hours and enumerated IFNγ-producing CD4+ T cells using intracellular staining and flow cytometry. We found that the CD4+ T cells in MHCII-expressing tumors produced strikingly greater amounts of IFNγ (Figure 3A, B), relative to their counterparts from MHCII-negative tumors at all time points tested, even after normalizing the numbers of T cells to tumor mass (Figure 3B). Despite this observed increase in cytokine production by CD4+ T cells in TS/A-hCIITA tumors relative to those in control tumors, CD4+ T cells in both groups experienced a progressive loss in the ability to produce IFNγ (Figure 3A, B), consistent with the idea that CD4+ T cells in both groups eventually became exhausted. Importantly, similar numbers of CD4+ T cells infiltrated MHCII-expressing and control tumors (Figure 3C). These results suggested that immune-suppressive mechanisms eventually overcome the immunologic advantage gained from tumor-specific MHCII expression.

**Figure 3.**
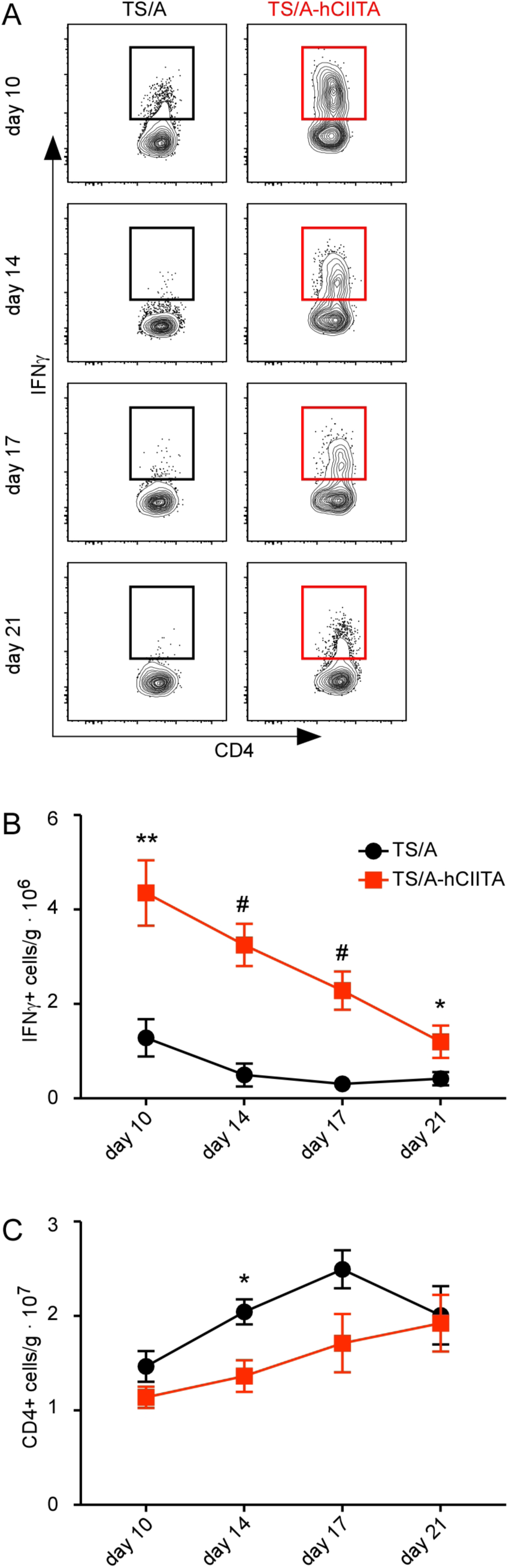
MHCII-expressing tumors promote CD4+ TIL activation. TS/A (black) and TS/A-hCIITA (red) tumors were excised on days 10, 14, 17, and 21 post-injection and single cell suspensions were restimulated in vitro. A. IFNγ production by CD4+ T cells was assayed by flow cytometry. Plots are gated on live, CD3+, CD4+ cells. B. The number of IFNγ-producing CD4+ T cells was normalized to tumor mass. C The total number of CD4+ T cells was normalized to tumor mass. This experiment contained 5 mice per group per timepoint and was performed two times with similar results. Error bars represent standard error of the mean. Statistical difference is expressed as *p<0.05, **p<0.005, ^#^p<0.0005.

### Tumor expression of MHCII enhances the CD8+ T cell response

CD4+ T cell help is required for effective cross-priming and antigen presentation to CD8+ T cells and for generating effective cytolytic responses (23, 28). Hence, we hypothesized that the presence of more activated CD4+ TILs in MHCII-expressing tumors would facilitate CD8+ T cell recruitment and activation, thereby enhancing their tumor-specific cytolytic capacity. To address this possibility, we injected TS/A and TS/A-hCIITA tumor cells into the mammary fat pad of BALB/c mice, harvested tumors at days 10, 14, 17, and 21 post injection and analyzed CD8+ TILs by flow cytometry. Given that TS/A tumor cells harbor an immunogenic murine leukemia virus (MuLV) (29), we initially enumerated CD8+ T cells that stained with a fluorescently-labeled MHCI tetramer containing a peptide from the MuLV env protein. We found similar frequencies of total and tumor-specific CD8+ T cells in both TS/A-CIITA and control tumors at early times (Figure 4A), but much higher frequencies of total and MuLV-specific CD8+ T cells in TS/A-hCIITA tumors on days 17 and 21. The higher frequencies of T cells were reflected in the numbers of total CD8+ T cells (Figure 4B) and the numbers of tumor-specific CD8+ T cells (Figure 4C), even when normalized to tumor mass.

**Figure 4.**
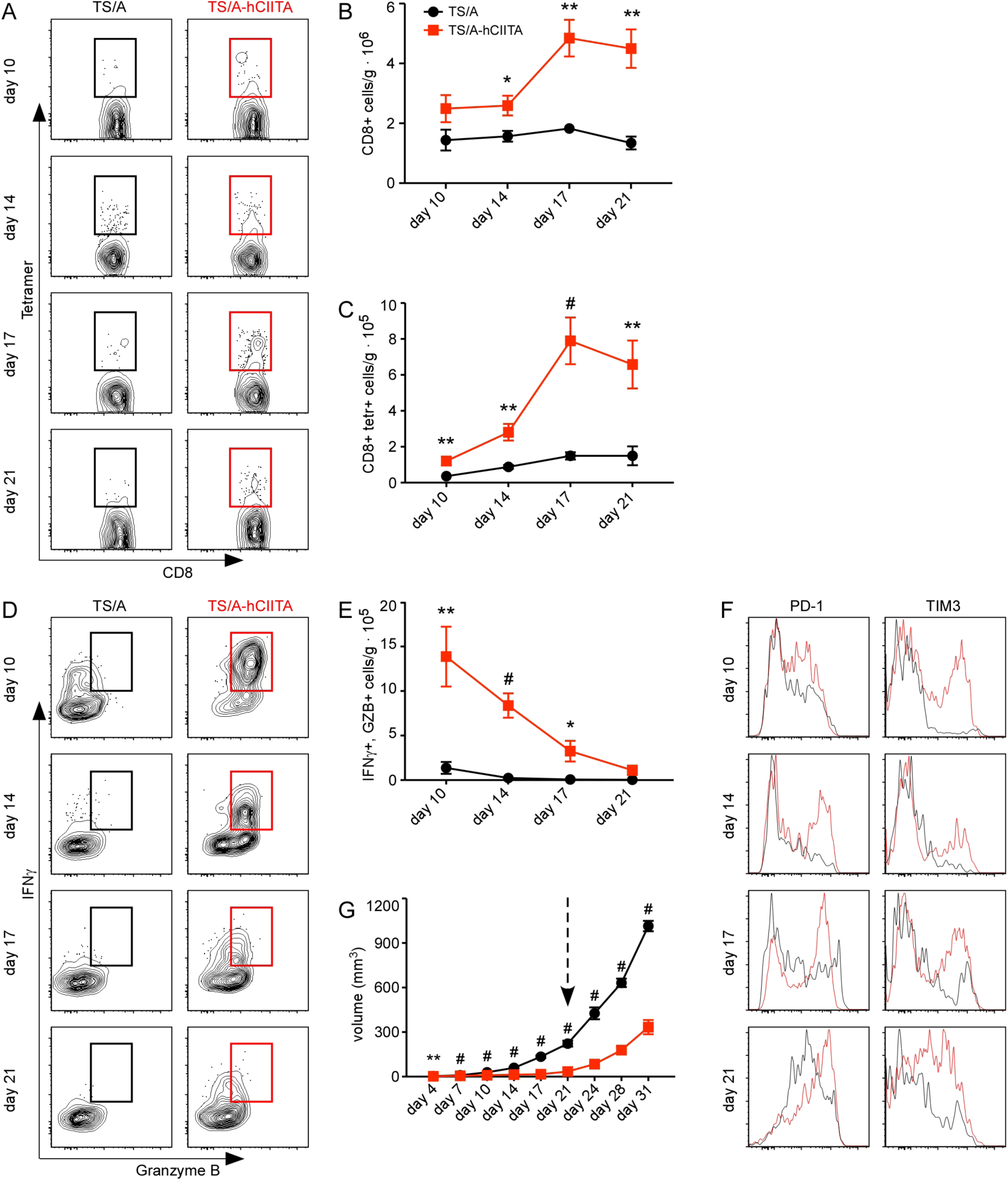
MHCII-expressing tumors maintain functional, tumor-specific CD8+ T cells. TS/A (black) and TS/A-hCIITA (red) cells were injected into the mammary fat pad of BALB/c mice, excised on days 10, 14, 17, and 21 post-injection and single cell suspensions were analyzed by flow cytometry. A. The frequency of MuLV env-specific CD8+ T cells was determined by tetramer binding and flow cytometry. Plots are gated on live, CD3+, CD8+ cells. B. The number of CD8+ T cells was normalized to tumor mass. C The number of MuLV env-specific CD8+ T cells was normalized to tumor mass. D. The ability of CD8+ T cells to make IFNγ and Granzyme B following restimulation ex vivo was determined by intracellular staining and flow cytometry. Plots are gated on live, CD3+, CD8+ cells. E. The number of CD8+IFNγ+GZB+ cells was normalized to tumor mass. F. The expression of PD-1 and TIM3 on CD8+ TILs was analyzed by flow cytometry. Histograms are gated on live, CD3+, CD8+ cells. G. The volume of TS/A and TS/A-hCIITA tumors was measured over an extended time frame and the arrow indicates a change in tumor growth rate corresponding to T cell exhaustion. These experiments contained 5 mice per group per timepoint and were performed at least two times. Error bars represent standard error of the mean. Statistical difference is expressed as *p<0.05, **p<0.005, ^#^p<0.0005.

We next assayed the functional ability of CD8+ T cells to produce IFNγ and granzyme B following restimulation *ex vivo*. We observed dramatically increased production of both IFNγ and granzyme B by CD8+ TILs from TS/A-hCIITA tumors relative to those in control tumors at all timepoints (Figure 4D). However, as we observed with CD4+ T cells, the CD8+ T cells in both groups progressively lost their ability to produce IFNγ and granzyme B (Figure 4D-E). Consistent with the idea that these cells might be exhausted, we observed that the expression of the inhibitory receptors, PD-1 and TIM3, are expressed by more CD8+ T cells from TS/A-hCIITA tumors than from control tumors at all time points. Moreover, the expression of these markers increased over time on CD8+ T cells (Figure 4F), even as T cell function declined.

Declining cytokine production and expression of inhibitory receptors is consistent with an exhausted phenotype, which occurs as early as day 10 in CD8+ T cells from control tumors and around day 21 in CD8+ T cells from MHCII-expressing tumors. To demonstrate the functional significance of CD8+ TIL exhaustion, we injected TS/A and TS/A-hCIITA cells into BALB/c mice and allowed tumors to grow for an extended period (Figure 4G). We found that although TS/A-hCIITA tumors initially grew more slowly than control tumors, they grew at similar rates following day 21 (Figure 4G), a time that coincides with the observed loss in T cell effector functions.

Given the expression of PD-1 on CD8+ T cells, particularly those in TS/A-hCIITA tumors, we hypothesized that PD-1 blockade would preserve CD8+ T cell function and further impair tumor growth. However, we found no difference in tumor growth following treatment with anti-PD-1 (Figure 5A). To investigate why PD-1 blockade had no impact on tumor growth, we assayed the expression of the PD-1 ligands, PD-L1 and PD-L2, on TS/A and TS/A-hCIITA tumor cells. We found that PD-L2 was constitutively expressed on the majority of CD45^neg^ cells, whereas PD-L1 was poorly expressed by CD45^neg^ cells (Figure 5B). Moreover, the over-expression of hCIITA had no impact on the expression of either ligand (Figure 5B). As a control for PD-L1 staining, we showed that PD-L1 was expressed at high levels by a portion of the CD45+ cells in both groups (Figure 5C). Thus, despite the exhaustion of tumor-infiltrating CD8+ T cells and high expression of PD-1, the blockade of PD-1 had minimal impact on T cell activity or tumor growth.

**Figure 5.**
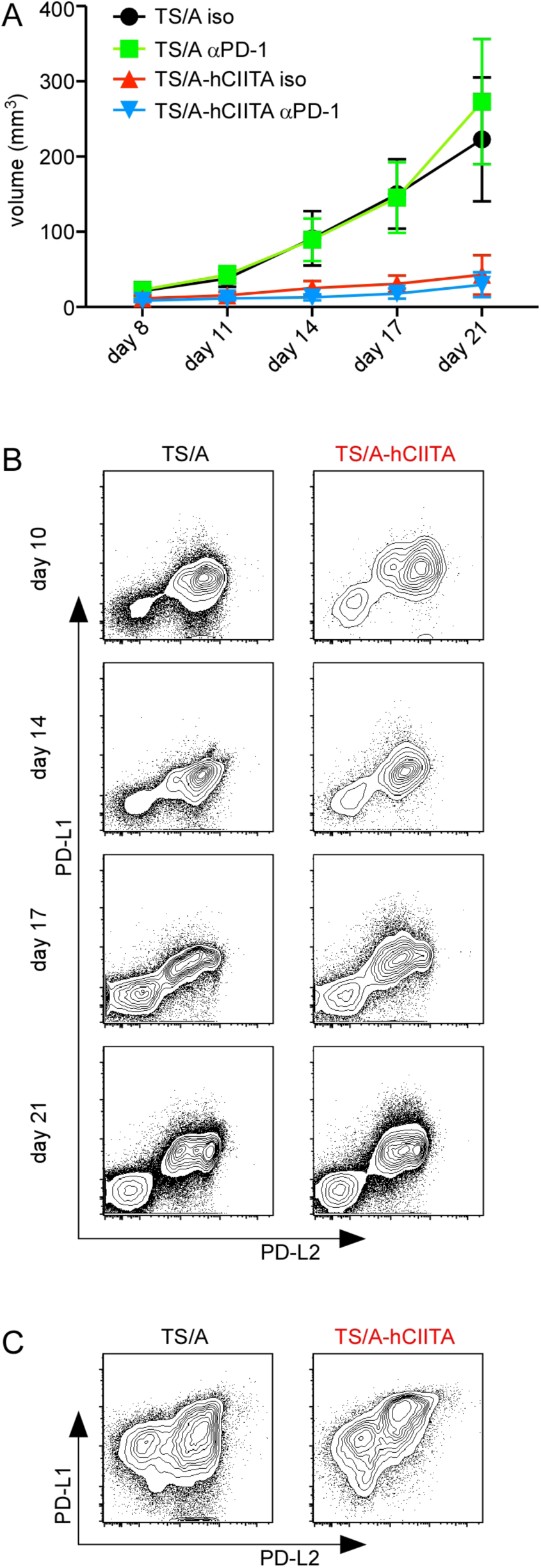
PD-1 blockade does not lead to tumor elimination. TS/A and TS/A-hCIITA cells were injected into the mammary fat pad of BALB/c mice and anti-PD-1 antibodies administered on days 8, 11, and 14 post injection. A. Tumor growth was monitored for 21 days. B-C. The expression of PD-L1 and PD-L2 on cell suspensions from disassociated tumors was analyzed by flow cytometry. Plots are gated on live, CD45^neg^ cells (B) or live, CD45+ cells (C). This experiment contained 5 mice per cohort per timepoint.

### CD4+ T cell depletion diminishes the benefit of MHCII expression

Our data suggested that local MHCII expression on tumor cells promoted the accumulation of activated CD4+ T cells, which indirectly supported CD8+ T cell accumulation and activation. To directly test this possibility, we depleted CD4+ cells prior to tumor cell implantation and evaluated tumor growth as well as the number and function of CD8+ T cells infiltrating TS/A-hCIITA and control tumors. As expected, we found that CD4 depletion restored the growth of MHCII-positive tumors to nearly that of MHCII-negative tumors in non-depleted mice (Figure 6A). However, we also found that CD4 depletion modestly increased the growth of MHCII-negative tumors, suggesting that CD4+ T cells have some role in promoting anti-tumor immunity via interactions with other, non-tumor, MHCII-expressing cells.

**Figure 6.**
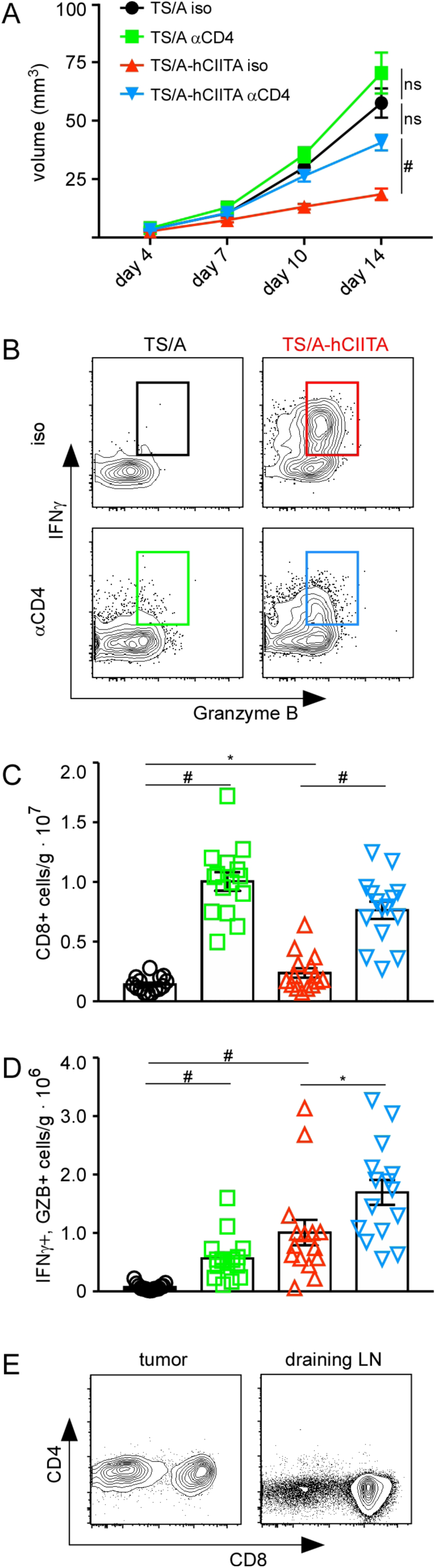
CD4 depletion impairs tumor growth and CD8+ T cell activity. BALB/c mice were treated with CD4-depleting or control antibodies on days −1 and +3 relative to the administration of TS/A and TS/A-hCIITA cells. A. Tumor volume was measured over 14 days. B. T cells from tumors were restimulated ex vivo and the expression of IFNγ and Granzyme B was assayed by flow cytometry on day 14. Plots shown are gated on live, CD3+, CD8+ lymphocytes. C. The number of CD8+ T cells was normalized to tumor mass. D The number of CD8+IFNγ+GZB+ cells was normalized to tumor mass. This experiment had 5 mice per group and was performed three times with similar results. Error bars represent standard error of the mean. Statistical difference is expressed as *p<0.05, **p<0.005, ^#^p<0.0005.

Importantly, CD4 depletion notably reduced the frequency of CD8+ T cells from MHCII-expressing tumors that produced IFNγ and granzyme B (Figure 6B), albeit not to the levels found in CD8+ T cells from MHCII-negative tumors. Surprisingly, the numbers of CD8+ T cells were dramatically increased in both MHCII-positive and MHCII-negative tumors following CD4 depletion (Figure 6C). Moreover, the numbers of CD8+ T cells producing IFNγ and granzyme B were also greatest in CD4-depleted TS/A-hCIITA tumors (Figure 6D). Absence of CD4+ staining in the tumor and draining lymph node confirmed elimination of CD4+ T cells following CD4 depletion (Figure 6E). These data suggest that despite the ability of CD4+ T cells to provide help to CD8+ T cells and facilitate the functional control of tumor growth, their absence does not entirely negate CD8+ T cell function and may even allow additional CD8+ T cell expansion via homeostatic mechanisms (30, 31).

### Expression of MHCII expands the responding T cell clones in tumors

The increased numbers of activated CD4+ and CD8+ T cells in MHCII-expressing tumors raised the possibilities that particular clones of T cells were expanding to a greater degree or that a broader array of T cell clones were being recruited to the tumor site. To test these possibilities, we harvested whole tumor and matched blood samples from three animals in each group on days 10, 14, 17, and 21 and sequenced the CDR3 region of the TCRβ chain to identify unique T cell clones. We first assessed the number of unique CDR3 reads in each sample, and found that estimated size of the TCRβ repertoire in the tumor was significantly smaller than that in peripheral blood (Figure 7A). Importantly, there was no difference in mean repertoire sizes between groups (Figure 7A), demonstrating that we have sufficient sequencing depth in each sample to appropriately compare them. We next compared the relative proportions of individual clones in each sample and found a relatively uniform distribution of (mostly infrequent) clones in blood, but found many examples of dramatically expanded clones in tumor samples (Figure 7B). Moreover, we observed more expanded clones and a greater magnitude thereof in TS/A-hCIITA tumors than in control tumors, whereas no differences were observed between blood samples from these same mice (Figure 7B). We quantified these results using the D50 repertoire diversity index of each cohort. We found that blood samples had high D50 scores that were not significantly different between TS/A-hCIITA and control tumor bearing mice, whereas tumor samples had low D50 scores and MHCII-expressing tumors were significantly lower than control tumors (Figure 7C). This result can be attributed to the local accumulation of the most abundant clonotypes, especially of the top 10 clones (data not shown). These data suggested that expanded T cell clones accumulated in tumors, but were minimally evident in the blood.

**Figure 7.**
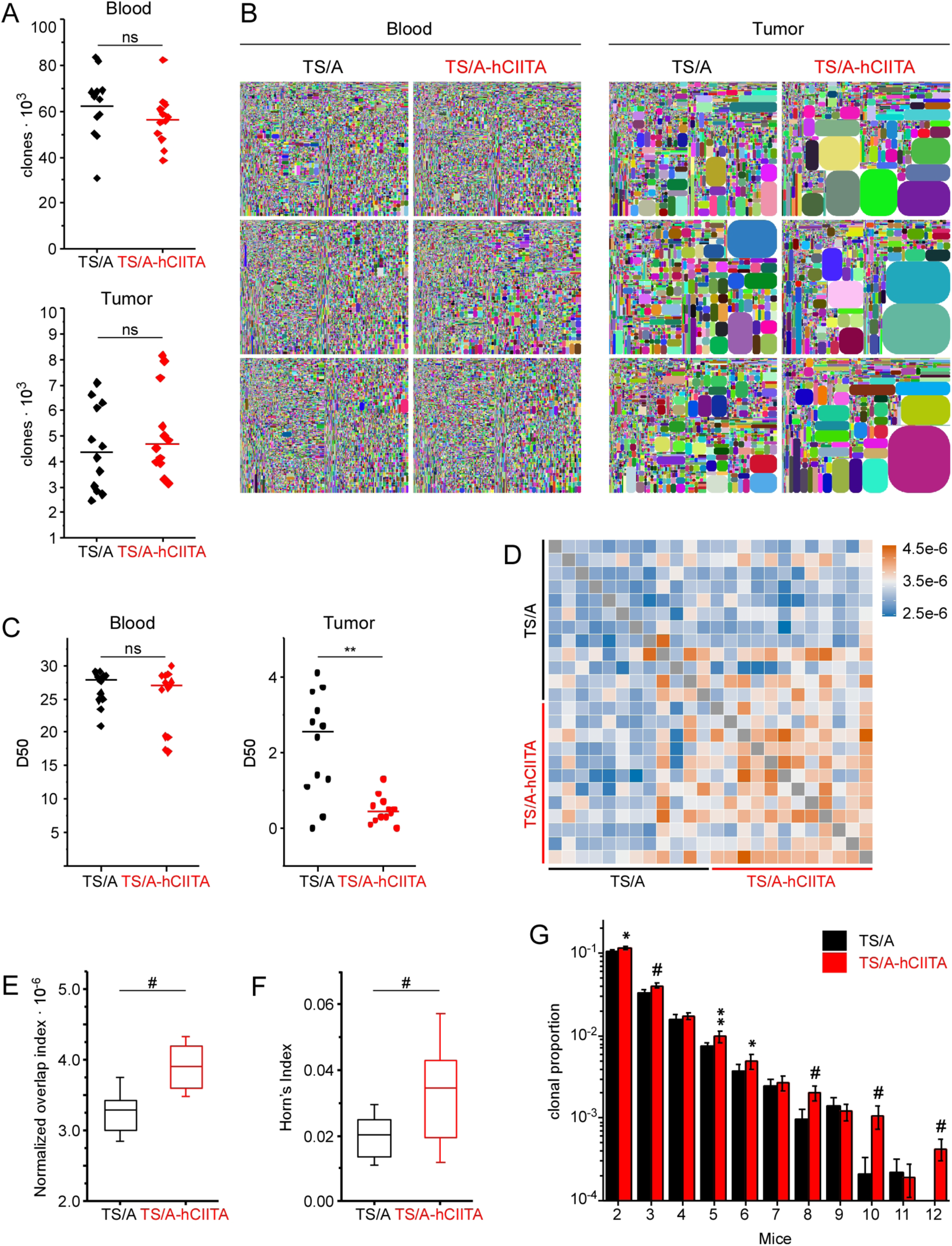
MHCII expression increases the number and magnitude of expanded TCR clonotypes in tumors, but not blood. TS/A and TS/A-hCIITA cells were injected and unique TCR clonotypes were determined by sequencing the TCRβ CDR3 regions in RNA purified from blood or tumors on days 10, 14, 17, and 21. A. The numbers of unique CDR3 reads were compared and horizontal lines denote group mean. B. Tree-map plots show relative sizes of individual clonotypes in matched blood and tumor samples on day 14. Each colored shape corresponds to a unique clone and its relative area indicates it representation in the repertoire. C. D50 repertoire diversity index quantifies expansion of detected clones in blood and tumor. D. Analysis of TIL CDR3 amino acid sequence similarity in all samples using pairwise overlap indexes, where legend values represent indexes. E. Quantification of pairwise comparison is demonstrated by plotting the overlap indexes. F. Overlap indexes were weighted by abundance of shared clones using Horn’s index. In E and F, error bars and horizontal lines indicate standard deviation and mean, respectively. G. Proportion of CDR3 amino acid sequences in each repertoire that are shared by a number (mice) of other repertoires in the same group (data represents 12 mice). Horizontal bars represent the mean and error bars are standard deviation. Statistical difference by t-test is expressed as *p<0.01, **p<0.001, ^#^p<0.0001

We next tested whether clonotypes of identical amino acid sequence were expanded in multiple mice, possibly because of dominant T cell responses to particular tumor antigens. By performing pairwise comparisons of normalized numbers of shared amino acid sequences, we found increased sharing of CDR3 sequences between TILs within MHCII-expressing tumors (Figure 7D). In addition, overlap indexes showed a significantly greater number of common sequences in TS/A-hCIITA tumors relative to controls (Figure 7E). Moreover, weighting sequence overlap by shared clones’ abundance using Horn’s pairwise overlap index also yielded a significant difference (Figure 7F). Finally, comparing sequences common to multiple mice (rather than two in pairwise comparison) revealed a significant increase in the number of shared sequences amongst TILs from MHCII-expressing tumors (Figure 7G). Collectively, these data suggested that MHCII expression on TS/A-hCIITA tumors increased intratumoral T cell clonal expansion, including clonotypes shared between mice, suggesting that they are responding to particular tumor antigens.

## DISCUSSION

Our data show that expression of MHCII on breast cancer cells enhances local CD4+ and CD8+ T cell responses and impairs tumor growth. CD4+ and CD8+ T cells that infiltrate MHCII-expressing tumors produce more effector molecules for longer periods, suggesting that locally-stimulated CD4+ T cells promote CD8+ T cell accumulation and function. Moreover, the overall breadth and magnitude of T cell clonal expansion is increased in MHCII-expressing tumors and similar clones of T cells are observed in multiple mice, suggesting that MHCII-expressing tumor cells likely present immunodominant epitopes that expand T cells with particular TCR rearrangements. Despite the local activation of both CD4+ and CD8+ T cells in MHCII-expressing tumors however, both cell types eventually become exhausted, allowing tumor outgrowth. These findings suggest that MHCII expression on tumor cells drives an intratumoral immune network that temporarily boosts local T cell responses and impairs tumor growth, but on its own cannot prevent T cell exhaustion.

MHCII molecules are typically expressed by professional antigen-presenting cells, like dendritic cells, which activate naïve CD4+ T cells in lymphoid organs. As a result, many tumors contain relatively few MHCII-expressing cells other than immunosuppressive cells of the myeloid lineage (32, 33). However, if tumor cells themselves express MHCII (and components of the MHCII antigen-processing pathway), then they should be able to directly stimulate CD4+ T cells. In fact, previous studies show that CIITA-mediated MHCII expression enables tumor cells to present intracellular antigens from various compartments (20, 34). Conceivably, this may allow CD4+ T cells to recognize previously unavailable antigens, then directly mediate anti-tumor functions and provide help to CD8+ T cells (35, 36), thereby enhancing immune control of tumor growth. Our data showing enhanced and prolonged CD4+ and CD8+ T cell function, expanded TCR clonotypes and reduced tumor growth are entirely consistent with this model.

Although MHCII-expressing tumor cells obviously present antigens to CD4+ T cells, it is unclear whether the enhanced functional activity of CD4+ T cells in MHC-expressing tumors is due to the re-activation of CD4+ effector T cells that were originally primed in the lymph node, or due to the direct priming and expansion of naïve CD4+ T cells by MHCII-expressing tumor cells. Dogma suggests that naïve T cells are most efficiently activated in lymph nodes; however, naïve T cells can be activated by non-traditional APCs in the absence of *bona fide* costimulatory signals if TCR signaling is sufficiently prolonged (37). Thus, if naïve CD4+ T cells do find their way into an MHCII-expressing tumor, they may be primed directly by tumor cells. Our data do not exclude this possibility, however we find numerous CD4+ T cells in MHCII-non-expressing tumors, suggesting that many tumor-infiltrating CD4+ T cells are initially primed by professional APCs and are subsequently re-activated by tumor cells in MHCII-expressing tumors.

Our data also show dramatically enhanced effector functions of CD8+ T cells in MHCII-expressing tumors, even though the expression of MHC class I molecules is unchanged, suggesting that locally activated CD4+ T cells produce factors capable of enhancing CD8+ T cell responses (36). Indeed, CD4 depletion largely negated the benefits of MHCII expression by tumor cells. Paradoxically, CD4 depletion also increased the number of CD8+ T cells in tumors, possibly due to homeostatic expansion (38). However, CD8+ T cell activation without help from CD4+ T cells can compromise their effector and memory functions (18, 39) and lead to exhaustion (21) - phenotypes that are consistent with the poor control of tumor growth that we observed following CD4+ T cell depletion.

The effect of local CD4+ T cell activation in MHCII-expressing tumors is also evident in the TCR repertoire, which has more and larger expansions of TCR clonotypes in MHCII-expressing tumors, but not in the blood of those same individuals. The expanded T cell repertoire in MHCII-expressing tumors may be due to epitope spreading (40), or to the local presentation of neoantigens or normally unavailable antigens as observed in melanoma patients (41).

Despite increased activation of both CD4+ and CD8+ T cells in MHCII-expressing tumors, both cell types eventually lose effector functions and become exhausted, allowing tumor outgrowth. This exhausted phenotype may be due to prolonged TCR signaling in the absence of inflammation or costimulation (42), a situation commonly encountered in tumors. In addition, tumors often acquire immune inhibitory characteristics. For example, although IFNγ is critical for anti-tumor immunity (20), prolonged IFNγ signaling often leads to “adaptive resistance”, in which tumor cells upregulate T cell-inhibitory proteins like indoleamine oxidase (IDO) or PD-L1 (43). As a result, the augmented IFNγ production by T cells responding to MHCII-expressing tumors may initially promote anti-tumor immunity, but later lead to immune suppression (44).

Although T cell exhaustion can sometimes be overcome with blocking antibodies to inhibitory receptors, like PD-1, in most cases, only a subset of patients respond to therapy. For example, patients whose melanoma tumors express MHCII are more likely to clinically benefit from PD-1/PD-L1 blockade (8). However, despite the increased expression of PD-1 on CD8+ T cells in our experiments, PD-1 blockade failed to restore T cell effector functions or promote tumor clearance, regardless of MHCII expression. This result is not entirely unexpected, as we failed to find substantial PD-L1 expression on tumor cells at any timepoint, a predictor of response to anti-PD-1 blockade (45). Moreover, T cell exhaustion is progressive and epigenetic restructuring can severely curtail the reinvigoration of T cells with PD-1 blockade (46). Hence, the efficacy of PD-1 blockade is mediated by numerous factors, including tumor phenotype, pre-treatment immune response and timing of administration.

Given the enhanced immune responses and better clinical outcomes in both mice and humans with MHCII-expressing tumors (8, 10, 13, 15)., how might we promote MHCII expression on tumors that currently lack MHCII expression? Therapeutic induction of MHCII on tumor cells should promote CD4+ T cell recognition of antigen, increase IFNγ production and enhance local CTL differentiation. The local administration of replication-deficient, oncolytic viruses that encode CIITA would likely promote MHCII expression and provide additional inflammatory signals via the activation of the anti-viral response (47, 48). An alternative mechanism for inducing MHCII expression *in vivo* is to alter the epigenetic control of gene expression in tumor cells using small molecules. Specifically, histone deacetylase inhibitors may directly induce tumor cell expression of MHCII (49) or enhance IFNγ-mediated CIITA expression (50). We would contend, however, that therapies triggering MHCII expression on tumor cells *in vivo* should be thought of as adjuvants for additional therapies that either promote anti-tumor immunity or are directly cytolytic to tumor cells.

In summary, our data show that MHCII expression on tumor cells enhances and prolongs both CD4+ and CD8+ T cell responses and slows tumor growth. However, T cells eventually become exhausted, leading to tumor outgrowth. Thus, combinatorial strategies are needed to enhance MHCII expression, prevent T cell exhaustion and promote tumor clearance.

## METHODS

### Cell culture, transfection, and selection

The TS/A murine mammary adenocarcinoma cell line was generously provided by Dr. Roberto S. Accolla, Department of Clinical and Biological Sciences, University of Insubria, Italy. TS/A cells were obtained at passage 22 and passaged 2 times prior to freezing. Cells were authenticated by assessing MHC haplotype via flow cytometry and by detection of antigens from murine leukemia virus. Transfected and control TS/A cells were also confirmed by gene expression of MHCII pathway gene products using nanostring assay. Frozen aliquots from passage 24 were thawed and expanded prior to injection. Cells were cultured in DMEM (Corning) supplemented with 10% fetal bovine serum (FBS; HyClone Laboratories, Inc.), and passaged by dissociating confluent monolayers with 0.05% trypsin, 0.53 mM EDTA without sodium bicarbonate (Corning), washing and reculturing. Cells were transfected with pcDNA3.1 with or without the hCIITA cDNA using the Xfect Transfection Reagent (Clontech) and selected in 500 µg/mL geneticin (Life Technologies). Clones that stably expressed MHCII were further selected by cell sorting. The expression of hCIITA, MHCII and CD74 in transfected cells was confirmed by Western blot using antibodies against hCIITA (E-12, Santa Cruz Biotechnology), MHCII (M5/114, Millipore), and CD74 (R&D Systems). Cells tested negative for mycoplasma (and 13 other mouse pathogens) via PCR performed by Charles River Research Animal Diagnostic Services on June 2015.

### RNA isolation and nanostring analysis

Total RNA was purified from excised tumors using the RNeasy Mini kit (Qiagen) and quantified by optical density at 260 nm using a DeNovix DS-11 spectrophotometer (DeNovix Inc.). Total RNA from blood was obtained from ∼1 mL (from mouse cardiac puncture) of whole blood collected into Paxgene RNA tubes using the Paxgene blood RNA kit (Qiagen) according to manufacturer instructions. The quality of RNA was assessed using Bioanalyzer 2100 (Agilent) and final concentration determined by Qubit (Life Technologies).

For nanostring analysis, 100 ng of purified RNA was added to 3 μL of Reporter CodeSet and 2 μL Capture ProbeSet in accordance with manufacturer’s recommendations (NanoString Technologies), and hybridized at 65°C overnight. Samples were processed the following day using the nCounter DxPrep Station and nCounter Dx Digital Analyzer (NanoString Technologies). Consumables were provided as an nCounter master kit by the manufacturer (NanoString Technologies). Data was normalized to housekeeping genes and analyzed using the nSolver 2.6 software. Heat maps were created using Cluster 3.0 and Java TreeView-1.1.6r4.

### Mice, tumor administration and antibody treatments

BALB/c mice of at least 6 weeks of age were purchased from Charles River Laboratories International, Inc. BALB/c.scid mice (CBySmn.CB17-*Prkdc^scid^I*J) were purchased from The Jackson Laboratory. Each experiment or time point represents cohorts of n=5 done at least in duplicate. Mice were injected with 1 × 10^5^ TS/A or TS/A-hCIITA tumor cells into the mammary fat pad on day 0. The longest (length, L) and shortest (width, W) tumor dimensions were measured by caliper and tumor volume was calculated using the formula, 0.4 x L x W^2^. For depletion experiments, 200 μg of CD4-depleting antibody (clone GK1.5, BioXCell) or CD25-depleting antibody (clone PC-61.5.3, BioXCell) or isotype-matched control antibody (clone LTF-2 or HRPN, respectively; BioXCell) were administered prior to tumor cell injection (day −1) and again 3 days following injection. For PD-1 blockade, 200 µg of anti-PD-1 antibody (clone J43, BioXCell) or isotype-matched control antibody (Armenian Hamster IgG, BioXCell) was administered on days 8, 11, and 14 post tumor cell injection. All procedures involving animals were approved by the University of Alabama at Birmingham Institutional Animal Care and Use Committee (IACUC) in protocol 09854.

### Tumor disassociation and T cell restimulation

Tumors were excised, weighed using an AL54 analytical balance (Mettler Toledo), and diced using a scalpel. Tumor fragments were then incubated in 2 mL RPMI1640 media (Lonza) supplemented with 5% FBS, 1.25 mg collagenase (c7657, Sigma) and 150 U DNase (d5025, Sigma), followed by gentle shaking at 200 rpm for 35 minutes at 37°C. Cell suspensions were filtered through 70 µm nylon cell strainers (Corning) and used for flow cytometry or T cell restimulation. For T cell stimulation, cells were washed and resuspended in RPMI1640, 5% FBS, 5 ng/mL phorbol 12-myristate 13-acetate (Sigma), 65 ng/mL ionomycin (ThermoFisher Scientific), and 10 µg/mL brefeldin A (Sigma) and cultured at 37°C for 5 hours.

### Flow cytometry and antibodies

Cell cycle analysis was performed by releasing non-confluent TS/A and TS/A-hCIITA cells from culture dishes using Accutase with 0.5 mM EDTA (Innovative Cell Technologies), fixing with 70% ethanol at 4°C overnight, and incubating them with 50 ug/ml propidium iodide (ThermoFisher Scientific) in PBS, 0.1% Triton-X100 (Sigma), and 50 ug/ml RNase at 37°C for 20 minutes in the dark. DNA content was determined using a BD FACSCalibur (BD Biosystems) flow cytometer and analyzed by ModFit LT version 3.3 (Verity Software House).

For immunophenotyping, cell suspensions from disassociated tumors were washed and resuspended in staining solution-PBS with 2% donor calf serum and 10 μg/ml FcBlock (2.4G2 -BioXCell)-for 10 min on ice before staining with fluorochrome-conjugated antibodies. Extracellular staining was done for 30 minutes, following by washing and fixation with 10% neutral buffered formalin solution (Sigma). Intracellular staining was performed on fixed cells permeablized with 0.1% IGEPAL CO-630 (Sigma) made in staining solution for 45 minutes. All staining was performed in the dark at 4°C. Antibodies against I-A/I-E (M5/114.15.2) and CD4 (GK1.5) were obtained from BioLegend). Antibodies against IFNγ (XMG1.2) and CD8 (53-6.7) were obtained from BD Biosciences. Antibodies against CD3 (17A2), KLRG1 (2F1), Granzyme B (NGZB), CD45.2 (104) and PD-1 (J43), were obtained from eBioscience. The MuLV class I tetramer-H2L^d^ (SPSYVYHQF) was obtained from the NIH Tetramer Core Facility at Emory University. The LIVE/DEAD red fixable dye was obtained from Life Technologies. Samples were run on a BDFACS Canto II system (BD Biosciences) data was analyzed using FlowJo version 9.9.

### T cell repertoire sequencing and analysis

For TCR library preparation, we used the commercially available iRepertoire platform (iRepertoire, Huntsville, AL) for nested amplicon arm-PCR of the CDR3 of the mouse TCR β-chains and addition of adaptors for Illumina platform sequencing. Reverse transcription of 300 to 500 ng of RNA was conducted with a One-Step reverse transcription and amplification kit (Qiagen) according to the manufacturer’s protocol. The PCR product was purified using Ampure XP magnetic beads (Agencourt), and secondary amplification of the resulting product was performed (TopTaq PCR Kit, Qiagen), allowing addition of Illumina adapter sequences (manufacturer’s protocol). Libraries were purified with Ampure XP magnetic beads and sequenced using Illumina MiSeq 150 nt paired-end read-length. The TCR CDR3 sequences were extracted from the raw sequencing data by iRepertoire. Briefly, raw paired-end fastq files were first demultiplexed based on the internal 6-nt barcode sequences added during library construction. The merged reads were mapped using a Smith-Waterman algorithm to germline V, D, J and C reference sequences downloaded from the IMGT web site (http://www.imgt.org). To define the CDR3 region, the position of CDR3 boundaries of reference sequences from the IMGT database were migrated onto reads through mapping results and the resulting CDR3 regions were extracted and translated into amino acids. Reading frames amino acid motif that was uninterrupted by a stop codon were identified as productive CDR3 amino acid sequences. Reads were filtered and error-corrected using iRepertoire proprietary SMART algorithm.

### Statistical Analysis

Hypothesis of difference testing between pooled replicate group means at each time point was done using multiple, unpaired independent t-tests with the Holm-Sidak method and assuming α=0.05. Statistical analyses were calculated using GraphPad Prism version 7.0a and all reported p-values are two-tailed. The TCR high-throughput sequencing data were analyzed in R environment using tcR package and common R routines. Diversity was measured using D50 immune repertoire diversity index. CDR3 sequence similarity between repertoires was assessed using overlap index and weighted Horn’s index available in the tcR package.

## Compliance with ethical standards

### Funding

This work was supported by the UAB Comprehensive Cancer Center (P30 CA013148) and by the Breast Cancer Research Foundation of Alabama.

## Ethical approval

All applicable international, national, and/or institutional guidelines for the care and use of animals were followed. All procedures performed in studies involving animals were in accordance with the ethical standards of the institution at which the studies were conducted. This article does not contain any studies with human participants performed by any of the authors.”

## Acknowledgements

The authors would like to thank Uma Mudunuru and Scott Simpler for animal husbandry, Eddy Yang MD, PhD and Debbie Della Manna of the UAB NanoString Laboratory and Enid Keyser of the UAB Comprehensive Flow Cytometry Core for lending respective expertise.

